# Alternative *c-MYC* mRNA transcripts as an additional tool for c-Myc2 and c-MycS production in BL60 tumors

**DOI:** 10.1101/2022.02.18.480896

**Authors:** Dina Ibrahim, Nathalie Faumont, Danielle Troutaud, Jean Feuillard, Mona Diab-Assaf, Ahmad Oulmouden

## Abstract

While studying c-Myc protein expression in several Burkitt lymphoma cell lines and in lymph nodes from a mouse model bearing a translocated *c-MYC* gene from the human BL line IARC-BL60, we surprisingly discovered a complex electrophoretic profile. Indeed, the BL60 cell line carrying the t(8;22) *c-MYC* translocation exhibits a simple pattern, with a single c-Myc2 isoform. Analysis of the *c-MYC* transcripts expressed by tumor lymph nodes in the mouse *λc-MYC (A*^*vy*^*/a)* showed for the first time five transcripts associated with t(8;22) *c-MYC* translocation. The five transcripts were correlated with the production of c-Myc2 and c-MycS, and loss of c-Myc1. The contribution of these transcripts to the oncogenic activation of the t(8;22) *c-MYC* is discussed.

## Introduction

The human *c-MYC* proto-oncogene is involved in the control of many cellular processes including cell growth and apoptosis (Thompson 1998). In normal cells, most transcripts of the human proto-oncogene *c-MYC* start at alternative promoters P1 and P2 and encode three proteins designated c-Myc1 (67 kDa), c-Myc2 (64 kDa) and c-MycS (55 kDa) (Stephen R. Hann et al. 1988) (Spotts et al. 1997) (Wierstra and Alves 2008). These three isoforms arise by alternative initiation of translation starting at three different in-frame codons: a non-canonical CUG for c-Myc1, an AUG located 15 codons downstream for c-Myc2 (Stephen R. Hann et al. 1988), and an AUG 100 codons further downstream for c-MycS (Spotts et al. 1997). The resulting proteins contain the same carboxy-terminal domain, but differ in their N-terminal regions. A third promoter (P3) within the first intron has also been described (Eick et al. 1990). P3 promoter transcriptional activity leads to an mRNA that lacks the N-terminal encoding sequence (15 amino acids) that is specific to c-Myc1. Therefore, the specific function of each of the c-Myc proteins lies in their N-terminal region. The ability to express at least three amino-terminally unique forms of c-Myc protein seems important for the normal function of c-Myc in cell growth control. Indeed, c-Myc1 and c-Myc2 possess different trans-activation efficiencies at the non-canonical CCAAT/enhancer-binding protein-binding site (S. R. Hann et al. 1994). Furthermore, c-Myc2 is predominant in growing cells while c-Myc1 is preferred as cells approach high-density growth arrest (Benassayag et al. 2005). In addition, c-Myc1 exhibits a strong induction of apoptosis compared to c-Myc2. c-MycS which is transiently expressed during rapid cell growth (Spotts et al. 1997), lacks the first 100 amino acids containing two phosphorylation sites (Thr58 and Ser62) involved in the stability of c-Myc proteins to proteosomal degradation (Escamilla-Powers and Sears 2007). Mutations of these sites are often linked to B-cell lymphomas and are correlated with reduced apoptotic potential (Benassayag et al. 2005). In normal mammalian cells, these c-Myc isoforms do not accumulate singly but in specific combinations or ratios characteristic of a given cell status. Therefore, imbalanced expression of the different c-Myc proteins directly contributes to the loss of cell growth control associated with tumor development (Benassayag et al. 2005).

In Burkitt’s lymphoma (BL) cells that have a *c-MYC* chromosomal translocation to one of the immunoglobulin (*IG*) loci on chromosomes 2, 14, or 22, the transcription of a *c-MYC* proto-oncogene is characterized by preferential transcription from the *c-MYC* promoter P1 (Gerbitz et al. 1999). Shifted promoter P2 to P1 leads to a change in the c-Myc1/c-Myc2 ratio that is often observed in human tumors cell lines (Dooley et al. 1994). In most cases, only c-Myc2 was detected with little or no c-Myc1. Several transgenic mouse models were generated to drive *c-MYC* expression throughout B-cell development under the control of different *Ig* enhancers, in order to mimic the *IG*-*c-MYC* translocation (Ferrad et al. 2020). The λc-MYC mouse model bearing a translocated *c-MYC* gene from the human BL line IARC-BL60 exhibits aggressive lymphomas with striking similarities to human BL (Kovalchuk et al. 2000). It was suggested that the mutation within the 5’ sequence and promoter shift from P2 to P1 leads to an unbalanced expression of the different c-Myc proteins in BL60. Indeed, BL60 exhibits an increased amount of c-Myc2 and fails to synthesize the detectable levels of c-Myc1 as assessed by Western blot (Cesarman et al. 1987) (Stephen R. Hann et al. 1988).

Here we report for the first time that tumors from lymph nodes of *λc-MYC (A*^*vy*^*/a)* mice exhibit five transcripts as new tools to exacerbate the imbalanced expression of c-Myc proteins that may explain the aggressive lymphomas in the *λc-MYC* (*A*^*vy*^*/a*) mouse model.

## MATERIALS AND METHODS

### Mouse Models

*λc-MYC(a/a)* mice on a C57BL/6 background were kindly provided by Pr. Georg BORNKAMM (Helmholtz Center, Munich, Germany). The *λc-MYC* (*A*^*vy*^*/a*) mouse model used in this work develops a human Burkitt-like lymphoma that is derived from crossing *λc-MYC (a/a)* mice with Yellow *A*^*vy*^*/a* mice. *A*^*vy*^ mice were kindly provided by Dr. David SKAAR (Department of Biological Sciences, Centre for Human Health and the Environment, North Carolina State,University, Raleigh, NC USA) for epigenetic studies. All animal experiments and protocols were conducted in accordance with European guidelines and regulations for animals used for scientific purposes, implemented in France as follows “Décret n°2012-118 du 1er février 2013 relatif à la protection des animaux utilisés à des fins scientifiques”. Considerable efforts were made to minimize the number of animals used and to ensure optimal conditions for their well-being and welfare before, during and after each experiment.

### Cell lines

Burkitt’s cell lines used in this work were obtained from the ATTC, cultured from cryopreserved cells, in RPMI-1640 medium supplemented with 10% fetal calf serum (FCS), 1% penicillin (100 µg/mL), 1% streptomycin (100 µg/mL), L-Glutamine, sodium pyruvate and vitamins. All cells were cultured in a humidified chamber at 37°C and 5% CO2. They were not stored beyond the 35th passage. The cell pellets were washed with PBS (phosphate-buffered saline) twice and then used for either RNA or protein extraction.

### RNA extraction

RNA was prepared using RNeasy Mini Kit (Qiagen, Les Ulis, France) according to the manufacturer’s instructions and treated with DNase I by using RNase free DNase set (Qiagen).

### Identification of 5’UTR, 3’UTR and full-length of cDNA

The SMARTer RACE 5’/3’ Kit (Takara Bio, Catalog no 634860, USA) was used to perform both 5’- and 3’-rapid amplification of cDNA ends (RACE) according to the manufacturer’s instructions. Briefly, 1µg of total RNA was used for (from lymph node tumors or cell lines) 5’-RACE-Ready cDNA and 3’ -RACE-Ready cDNA. Primer pairs P5’UTR1/UPM (Universal Primer A Mix) and P5’UTR2/UPM were used for PCR amplification using 5’-RACE-Ready cDNA as template to obtain 5’UTR, P3’UTR1/UPM was used to obtain 3’UTR by PCR amplification using 3’ -RACE-Ready cDNA as template. By cloning and sequencing (BigDye™ Direct Cycle Sequencing Kit - Thermo Fisher) the 5’ and 3’ UTRs we designed specific primers to obtain full-cDNA and the corresponding mRNA structure. All primers used in this work are listed in Table1. PCR amplification was performed as follows: denaturation 2 min at 94°C, 35 cycles (94 °C 15 sec, 61 °C 15 sec, 72 °C 1 min), 72°C 2 min.

**Table 1:**
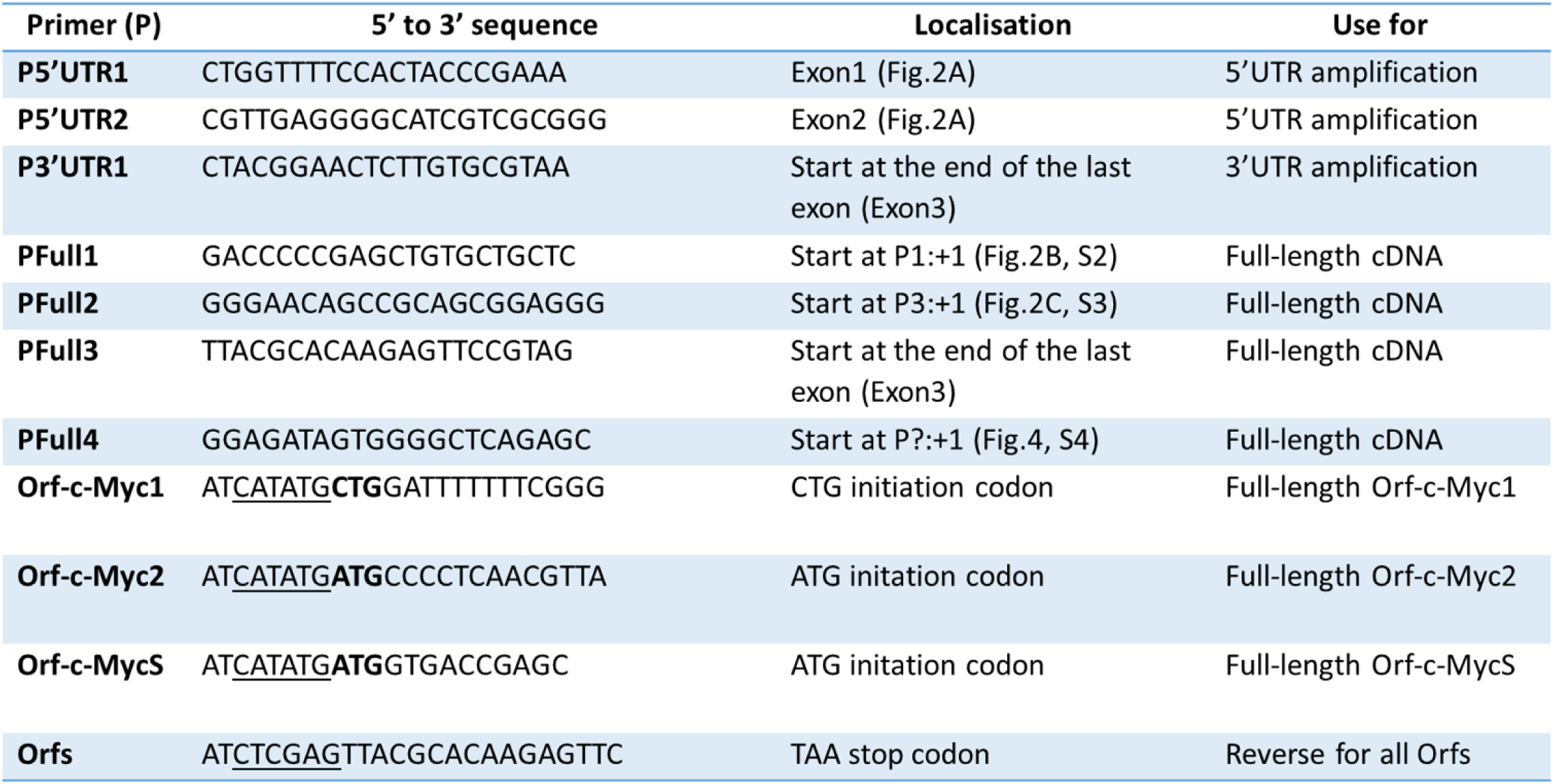
Primers (P) used to obtain 5’UTR, 3’UTR and total length cDNAs of c-MYC transcripts (only open reading frames (ORFs) are indicated). The underlined sequences of Orf-c-Myc1, Orf-c-Myc2 and Orf-c-MycS corresponds to NdeI restriction sites; the underlined sequence of Orfs corresponds to XhoI restriction site; **CTG** and **ATG**: Translation start codons.

### *In vitro* expression of c-Myc1, c-Myc2 and c-MycS reading frames

Reading frame sequences encoding each protein were amplified by PCR (as described above) using primer pairs described in Table 1 and cloned in the pT7CFE1 expression vector (Figure S1). *In vitro* transcription and translation were performed according to manufacturer’s instruction (pT7CFE1-CHis Vector for Mammalian Cell-Free Protein Expression; Thermo Fisher).

### Protein extraction and Western blot

Fresh or frozen animal tissues (30 mg) were dissected on ice. 2×10^6^ cells (extracted tissues or cell culture) were suspended in 50 μL RIPA (Radio ImmunoPrecipitaion Assay; BioRad) lysis buffer containing 200 mM PMSF (Alpha-Phenyl Methyl Sulfonyl Fluoride, serine protease inhibitor), 100 mM sodium orthovarnadate, and a protease inhibitor cocktail (Santa-Cruz Biotech). Lysis was performed on ice for 30 minutes. The soluble protein fraction was recovered after centrifugation at 13,000 rpm at 4°C for 15 minutes. Proteins were assayed by the Bradford method. Then proteins extracted or *in vitro* synthesized proteins, were denatured at 95°C for 3-5 minutes in the presence of β-mercaptoethanol and Laemmli blue.

Equal amounts of denaturated proteins (30 µg/lane) were separated by SDS-PAGE then transferred to PVDF membrane. Nonspecific binding sites were blocked for 1 h with 5% non-fat dry milk in TBS containing 0.1% Tween-20. After overnight incubation at 4°C with specific primary Ab (9E10, dilution 1/200, sc-40 Santa Cruz Biotechnology), membranes were incubated with appropriate HRP-conjugated secondary Ab (dilution 1/1000, sc-2357 Santa Cruz Biotechnology) for 1 h at room temperature and revealed by an enhanced chemiluminescent detection method (Immubilon Western, Millipore). Protein-loading control was performed with GAPDH or β-actin Ab.

## Results

### *c-MYC* gene translocation and synthesis of c-Myc proteins in Burkitt lymphomas

In our *λc-MYC (A*^*vy*^*/a)* model mouse (Figure 1) of Burkitt lymphomas (Kovalchuk et al. 2000), we repeatedly obtained a complex electrophoretic pattern on Western blot, notably in lymph node tumors. We consistently observed two bands that could correspond to c-Myc2 and c-MycS with an apparent molecular weight of 64 and 55 kDa respectively in those tumors (Figure 2A). All Burkitt lymphoma cell lines studied exhibit doublet bands (Figure 2B): c-Myc2 (64 kDa) and probably its phosphorylated (Stephen R. Hann et al. 1988) form (65 kDa). Burkitt lymphoma cell lines are considered to express only c-Myc2 (Stephen R. Hann et al. 1988). Since we were unable to explain the presence of the other bands (Figures 2A, 2B), we performed a control test to detect c-Myc proteins.

**Figure 1:**
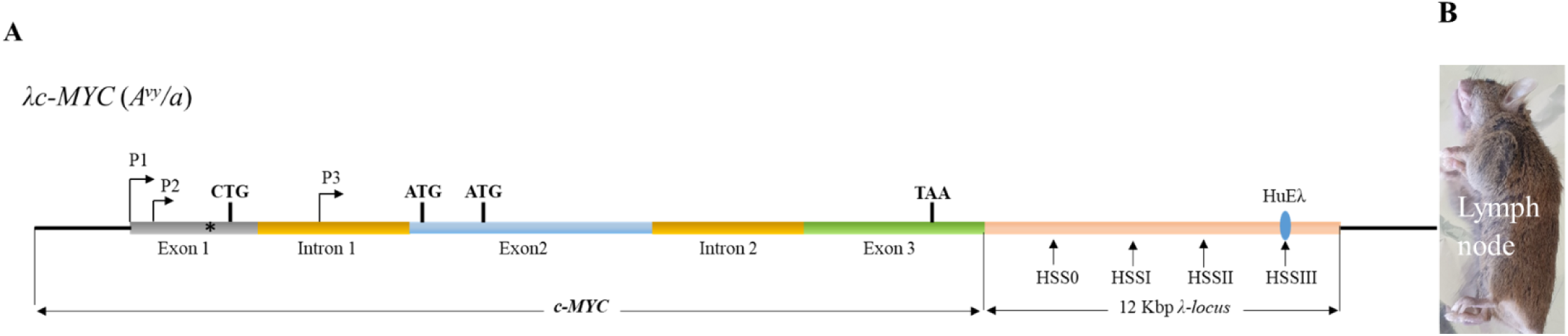
Structure of the λc-MYC transgene (A) carried by the λc-MYC (A^vy^/a) model mouse of Burkitt’s lymphoma (B) (Kovalchuk et al. 2000). Diagram is not to scale.

**Figure 2:**
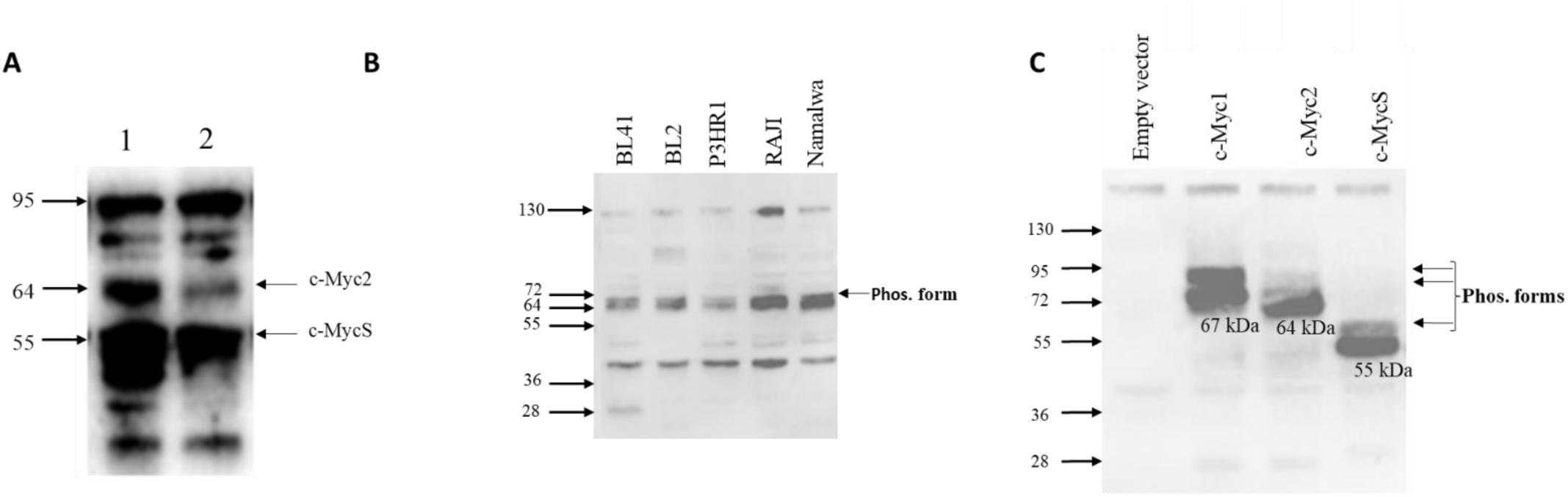
Analysis of the migration profile of c-Myc proteins by Western blot using the monoclonal antibody 9E10 directed against the conserved C-terminal part of c-Myc1, c-Myc2 and c-MycS. (A) Typical result obtained from cell extracts prepared from lymph nodes from two λc-MYC (A^vy^/a) mice (lane 1, lane 2). (B) Result obtained from extracts prepared from BL cell lines.(C) Electrophoretic pattern of c-Myc1, c-Myc2 and c-MycS proteins, obtained in vitro by transcription and translation of the open reading frames (Orf-c-Myc1, Orf-c-Myc2 and Orf-c-MycS) cloned separately into the pT7CFE1 vector (Figure S1). The apparent molecular weight of the different bands is shown on the left; Empty vector: negative control; Phos. form: Phosphorylated form

### c-Myc protein production by cell-free translation of a single cloned reading frame

The reading frame encoding c-Myc1, c-Myc2 or c-MycS was cloned into the pT7CFE1 vector (Figure S1), and the corresponding mRNA was translated *in vitro* using a mammalian *in vitro* translation system based on HeLa cell lysates. This experimental approach yielded the proteins detected in Figure 2C. In each case, significant band with apparent molecular weights of 67 kDa (c-Myc1), 64 kDa (c-Myc2) and 55 kDa (c-MycS) were obtained. A minor band corresponding to the phosphorylated forms (Stephen R. Hann et al. 1988) was also detected. However c-MycS lacks the first 100 amino acids, which contain two phosphorylation sites. Moreover, other phosphorylation sites have been described in the C-terminal region of the c-Myc proteins (Wasylishen et al. 2013). This result validates the capability of the 9E10 antibody used against the C-terminal region to detect the three isoforms of c-Myc protein, and our hypothesis with regard to the bands described in Figures 2A and 2B. Of note, c-Myc1 (P67) was undetectable in both lymph nodes (this work) and Burkitt’s lymphoma cell lines (this work and (Stephen R. Hann et al. 1988)).

### *c-MYC* transcripts in lymph nodes from *λc-MYC* mice

The *λc-MYC* mouse model was obtained by transgenesis using a 12 Kbp DNA fragment (Figure 1A) from the *c-MYC* translocation t(8;22) of BL60 cell line (Gerbitz et al. 1999) (Kovalchuk et al. 2000). As the electrophoretic profile of c-Myc proteins in tumors from lymph nodes from *λc-MYC (A*^*vy*^*/a)* mice was no longer the similar to that described for the BL60 line, with c-Myc2 but also c-MycS (apparent molecular weight of 64 and 55 kDa respectively, Figure 2A), we then analyzed the *c-MYC* transcripts in these tumors.

Total RNA was extracted and the 5’ and 3’ regions were obtained by RACE-PCR. The *c-MYC* specific primers used are described in Table1. To obtain the 5’UTR (5’UnTranslated Region), we used the primers P5’UTR1 and P5’UTR2 located at exon 1 and exon 2 respectively (Figure 3A) to account for the different promoter activities (P1, P2 or P3) described in the literature. The primer P3’UTR1 (Figure 3A) was used to obtain 3’UTR. The 5’ and 3’ established regions allowed us to define PFull1, PFull2 and PFull3 primers (Figures 3B, 3C) to obtain the total cDNA and the structure of *c-MYC* mRNAs after sequencing. For the 3’ region we repeatedly obtained (Figure S2), the same 3’UTRs for both the lymph nodes and for the BL41 cell line corresponding to the use of the two described polyadenylation sites (Laird-Offringa et al. 1989).

**Figure 3:**
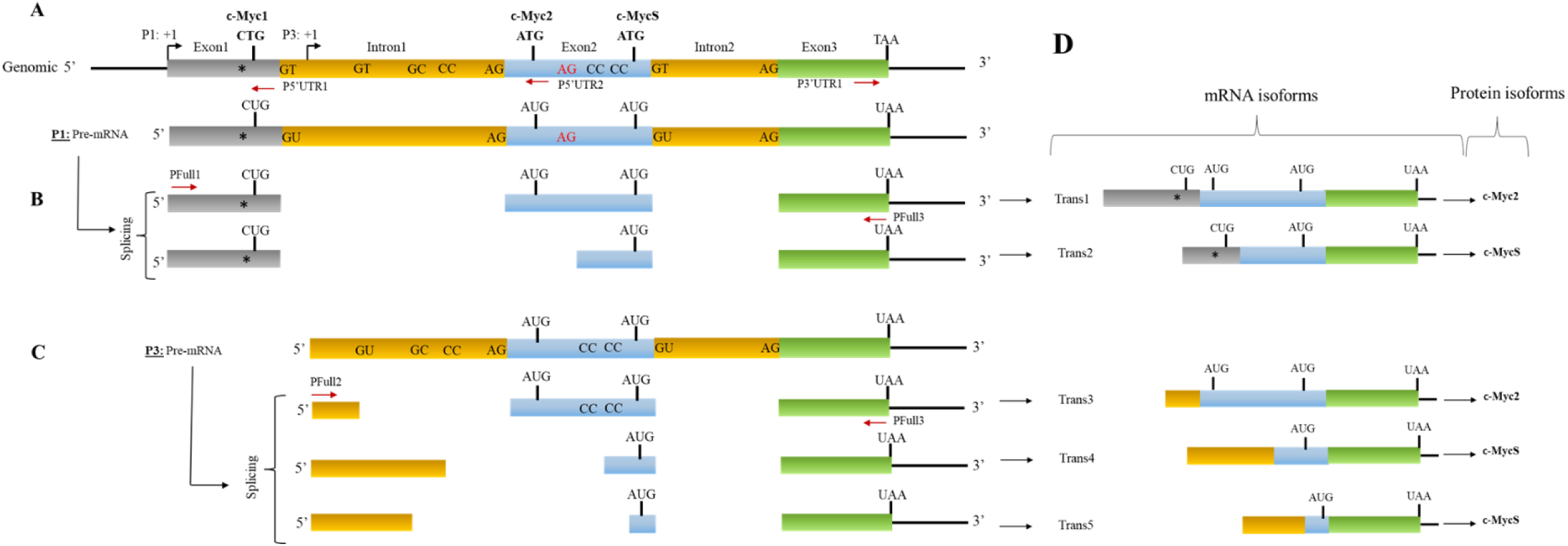
Different events leading to the genesis of several mRNAs of the c-MYC gene related to genomic structure. Genomic structure (A) at the origin of the different events (B), (C) and (D). Primers P5’UTR1 and P5’UTR2 used to obtain the 5’UTR; primer P3’UTR1 to obtain the 3’UTR and the other primers were used as indicated to obtain the full-length cDNA. The dinucleotides indicated represent the different splice donor and acceptor sites. The triplets indicate the translation initiation codons to obtain c-Myc1, c-Myc2 or c-MycS. TAA: Translation stop codon. *: PvuII mutation. Diagram is not to scale.

Sequence analysis of the cDNA revealed the presence of five *c-MYC* transcripts (Figure 3D), two from transcription through P1 (P1 promoter) and three from P3 (P3 promoter) which is located at intron 1 (Figure 3A). Transcription from P1 has been reported in the BL60 line, but to our knowledge, the nature of the transcripts has not been described. Transcription from P3 has not been reported in the BL60 cells. It is noteworthy that Trans1 (transcript 1) carries the PvuII mutation which has been suggested to be responsible for the production of the single c-Myc2 variant in BL60 cells (Figure 3D and S3). All *c-MYC* transcripts described in this work differ at the 5’UTR and/or at exon 2 (Figure 3). They all have the entire last exon (exon3). In addition the 5’UTRs of three transcripts (Trans1,2,3) contains termination codons in all three reading frames whereas the two 5’UTR transcripts (Trans4,5) (Figure S3) are in reading frames of a part of exon 2 and the coding region of c-MycS (see discussion).

### P1 promoter transcripts and c-Myc proteins

The genomic region corresponding to transcripts 1 and 2 (Trans1 and Trans2) resulting from P1 promoter use (Figures 3 and S2) revealed that the mature Trans1 mRNA (Figures 3D and S2) is derived from the canonical splicing of a *c-MYC* pre-mRNA (Figures 3B and S2) previously described (Watt et al. 1983). However, Trans2 (revealed in this work) is obtained by an alternative splicing acceptor site within exon 2 (Figures 2B and S3). In both cases, the donor (GT) and acceptor (AG) sites are canonicals. Translation of the two transcripts in the three reading frames predicts (Figure 3D) the formation of c-Myc1 (Trans1) and c-MycS (Trans2) isoforms. Nevertheless, our Western blot experiments (Figure 2A) do not detect the c-Myc1 (p67) isoform. This result is in agreement with previously reported data showing that a substitution near the non-canonical CUG initiation site prevents translation from the CUG site and thus c-Myc1 production (Stephen R. Hann et al. 1988). It is therefore very likely that the translation of Trans1 leads to the formation of c-Myc2 from the AUG initiation codon located at exon 2 (Figures 3 and S3). Taken together, analysis of the two transcripts coupled with Western blot data (Figure 2A) suggest production of c-Myc2 (instead of c-Myc1) and c-MycS by *in vivo* translation of Trans1 and Trans2 respectively.

### P3 promoter transcripts and c-Myc proteins

A similar analysis of the P3 promoter transcripts (Trans3, Trans4 and Trans5) revealed (Figures 3C and S4) that the Trans4 transcript is derived from the splicing of a pre-mRNA with canonical donor and acceptor sites located at intron 1 while respecting the canonical sites between exon 2 and exon 3. Since Trans4 lacks exon 1 (Figure 3C), translation of this mRNA *in vivo* will necessarily lead to the production of c-Myc2. The two other transcripts (Trans3 and Trans4) were derived from alternative splicing of pre-mRNA using different non-canonical splice donor and acceptor sites at intron 1 and exon 2. If these mRNA were translated *in vivo* they would produce at least (see discussion) the c-MycS isoform. Overall, P3 transcripts will result in the enrichment of c-Myc2 and c-MycS isoforms.

### *c-MYC* transcripts and c-Myc proteins in the BL41 cell line

Data described above led us to carry out a study of *c-MYC* transcripts and the corresponding proteins in a Burkitt lymphoma cell line.

Total RNA were extracted and subjected to the same RACE-PCR approach to obtain the 5’UTR region and then the total *c-MYC* mRNA expressed by BL41 cells. We used the same primers as above to obtain the 5’ and 3’ UTRs. We obtained amplifications with primer P5’UTR2 located in exon 2 (for 5’UTR) and primer P3’UTR1 located in exon 3 (for 3’UTR). We did not obtain any amplification with the other primer P5’UTR1 located in exon 1.

Sequence analysis of cDNA corresponding to the mRNA showed a single repeatedly obtained transcript (Figures 4 and S5). The 5’UTR was formed by a hybrid sequence including an *IGH* region and the end of intron 1 of *c-MYC*. The rest of the mRNA was transcribed from exons 2 and 3. This hybrid transcript was probably the result of transcriptional activity at the *IGH* locus following the t(8;14) translocation that characterizes the BL41 cell line. The fact that we did not obtain any amplification with the P5’UTR1 primer supports the presence of only *c-MYC* transcripts lacking exon 1. The translation of this mRNA predicts c-Myc2 protein in agreement with previously described data and with the results obtained by Western blot (Figure 2B).

**Figure 4:**
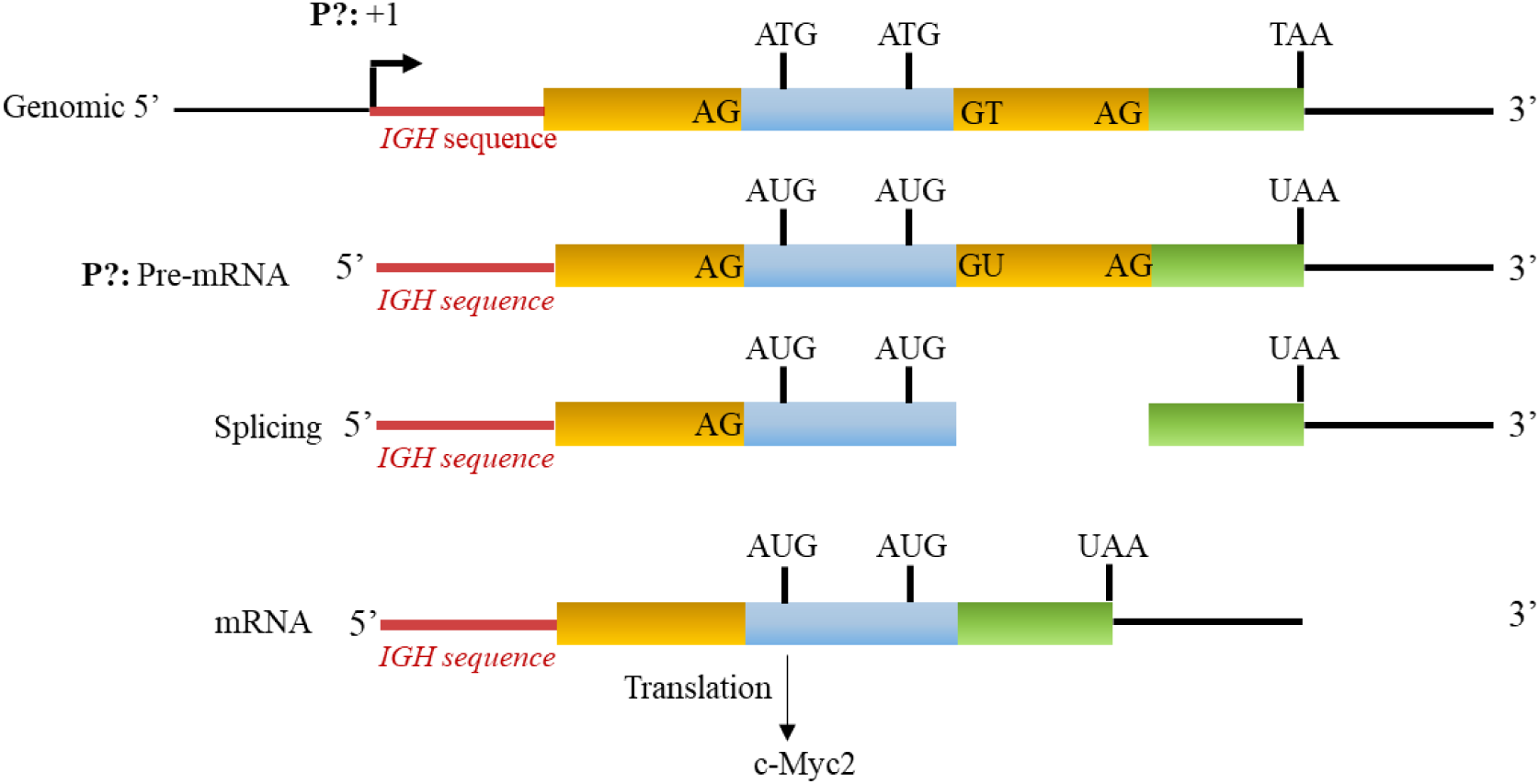
Partial genomic structure of the c-MYC gene associated with t(8;14) translocation of BL41. The expression from a yet to be determined promoter (P?) generates a hybrid mRNA with an IGH sequence. The translation of the resulting mRNA leads to the production of the c-Myc2 protein. Diagram is not to scale.

## Discussion

In this study, we report for the first time that tumor lymph nodes from the *λc-MYC (A*^*vy*^*/a)* mice exhibit five different *c-MYC* mRNA that invariably lead to the production of c-Myc2 and/or c-MycS shown on Western blot.

It is now well established that the quantitative and qualitative expression of the two proteins c-Myc1 and c-Myc2 and to a lesser extent c-MycS play an important role in the control of normal (non-cancerous) cell proliferation and cell homeostasis (S. R. Hann et al. 1994) (Batsché and Crémisi 1999). Although the ratio between these three isoforms is difficult to determine because their quantity changes according to stages of the cell cycle, the mechanisms that support the production of c-Myc2 and/or c-MycS promote a cancerous state: the *c-MYC* proto-oncogene becomes the *c-MYC* oncogene (Hann et. 1999) (Spotts et al. 1997).

Virtually all Burkitt lymphoma cell lines show a rearrangement between the *c-MYC* locus and one of the immunoglobulin gene loci. These rearrangements alone or in combination with mutations in the 5’ region of the *c-MYC* gene often lead to the production of c-Myc2 alone. Previously, it was shown that the BL60 line carrying the t(8;22) translocation exhibits a shift from promoter 2 to promoter 1 and harbors a substitution that prevents translation from the upstream non-canonical CUG translation site. Thus, BL60 was described to have lost c-Myc1 production and to produce only c-Myc2 (S. R. Hann et al. 1994). It should be noted that the 12 kbp DNA fragment (Figure 1A) from the t(8;22) translocation of the BL60 line used to obtain the λc-MYC mouse (Kovalchuk et al. 2000) contains all the genetic information (including the *c-MYC* gene) necessary for the development of an aggressive B lymphoma, particularly in the lymph nodes. In agreement with these previous data, the Trans1 mRNA discovered in this work was derived from the P1 promoter (shift from P2 to P1, as in BL60) and carries the described substitution near the CUG translation site as reported for translation of the only open reading frame of Trans1 which codes for the the production c-Myc1. Nevertheless, its presence in Western blots remains undetectable. This observation supports the fact that a mutation near the CUG translation site prevented its use as a translation initiation site and strongly suggests that *in vivo* translation of Trans1 leads to the formation of c-Myc2.

The second mRNA (Trans2) that we found is also from the P1 promoter, also carries the substitution near CUG, but with alternative splicing at exon 2. The only Myc protein that will be produced if this Trans2 is translated *in vivo* will be c-MycS. This hypothesis is in agreement with our Western blot data.

The other three transcripts (Trans3, 4 and 5) were derived from transcription at the P3 promoter located in intron 1. The translation of Trans3 predicts the production of c-Myc2 according to the only open reading frame. However, the predictions that can be drawn from the primary structure of Trans4 and 5 are less clear. Indeed, despite different alternative splicing, Trans4 and 5 each present an open reading frame from, respectively, the first and second nucleotides of transcription initiation to the stop codon at the last exon 3 of the *c-MYC* gene. Several potential non-AUG (Diaz de Arce, Noderer, and Wang 2018) sites were located in the 5’ region of these two mRNA but their use as translation initiation sites remains to be determined experimentally.

Alternative splicing of pre-mRNA and/or alternative use of promoters play an important role in the genesis and development of different types of cancer. Indeed, many cases of splicing isoforms that lead or promote cancer progression have been identified. For example, cyclin D1b, the oncogenic form of cyclin D1, expressed in lymphomas, results from alternative splicing. Another example is the anti-apoptotic isoform Bcl-xL which is strongly expressed in a large number of tumors to ensure high cell survival (Paronetto, Passacantilli, and Sette 2016). Finally, the use of alternative promoters and/or alternative splicing generates a complex pattern of pro- and anti-apoptotic isoforms of the TP73 gene (Zaika et al. 2002) which is strikingly similar to the mechanism used by the t(8;22) translocation as found in this study, to generate only oncogenic forms of c-Myc.

In summary, we report for the first time the concept of multi-transcript genesis as a tool for the development of Burkitt’s lymphoma in the case of t(8;22) translocation. Our work suggests that this type of approach may be useful for the identification of *c-MYC* transcripts with B lymphoma. Of course, these transcripts will differ depending on the type of B lymphoma associated with specific *c-MYC* dysregulation. The BL41 line used in this work is an example.

In this context, an exhaustive study of *c-MYC* transcripts becomes necessary. A nanopore sequencing approach is currently underway and data obtained will provide new diagnostic tools.

## Acknowledgements

We would like to thank Dr J Cook Moreau, UMR CNRS 7276, Limoges, France, for the English revision. This study was supported by grants from the Institut National du Cancer (INCa), the Ligue Contre le Cancer (Equipe Labellisée Ligue), the Comité Orientation Recherche Cancer (CORC), the Limousin Region and the Haute Vienne and Corrèze committees of the Ligue Nationale contre le Cancer and by the Lyons Club de Corrèze.

## References

Batsché, E., and C. Crémisi. 1999. “Opposite Transcriptional Activity between the Wild Type C-Myc Gene Coding for c-Myc1 and c-Myc2 Proteins and c-Myc1 and c-Myc2 Separately.” Oncogene 18 (41): 5662–71. https://doi.org/10.1038/sj.onc.1202927.

Benassayag, C., L. Montero, N. Colombié, P. Gallant, D. Cribbs, and D. Morello. 2005. “Human C-Myc Isoforms Differentially Regulate Cell Growth and Apoptosis in Drosophila Melanogaster.” Molecular and Cellular Biology 25 (22): 9897–9909. https://doi.org/10.1128/MCB.25.22.9897-9909.2005.

Cesarman, E., R. Dalla-Favera, D. Bentley, and M. Groudine. 1987. “Mutations in the First Exon Are Associated with Altered Transcription of C-Myc in Burkitt Lymphoma.” Science (New York, N.Y.) 238 (4831): 1272–75. https://doi.org/10.1126/science.3685977.

Diaz de Arce, Alexander J., William L. Noderer, and Clifford L. Wang. 2018. “Complete Motif Analysis of Sequence Requirements for Translation Initiation at Non-AUG Start Codons.” Nucleic Acids Research 46 (2): 985–94. https://doi.org/10.1093/nar/gkx1114.

Dooley, S., I. Wundrack, C. Welter, and N. Blin. 1994. “Constitutive C-Myc Overexpression and P1/P2 Promoter Shift in a Small-Cell Lung-Cancer Cell-Line.” International Journal of Oncology 5 (1): 65–68.

Eick, D., A. Polack, E. Kofler, G. M. Lenoir, A. B. Rickinson, and G. W. Bornkamm. 1990. “Expression of P0-and P3-RNA from the Normal and Translocated c-Myc Allele in Burkitt’s Lymphoma Cells.” Oncogene 5 (9): 1397–1402.

Escamilla-Powers, Julienne R., and Rosalie C. Sears. 2007. “A Conserved Pathway That Controls C-Myc Protein Stability through Opposing Phosphorylation Events Occurs in Yeast.” The Journal of Biological Chemistry 282 (8): 5432–42. https://doi.org/10.1074/jbc.M611437200.

Ferrad, Melissa, Nour Ghazzaui, Hussein Issaoui, Jeanne Cook-Moreau, and Yves Denizot. 2020. “Mouse Models of C-Myc Deregulation Driven by IgH Locus Enhancers as Models of B-Cell Lymphomagenesis.” Frontiers in Immunology 11: 1564. https://doi.org/10.3389/fimmu.2020.01564.

Gerbitz, A., J. Mautner, C. Geltinger, K. Hörtnagel, B. Christoph, H. Asenbauer, G. Klobeck, A. Polack, and G. W. Bornkamm. 1999. “Deregulation of the Proto-Oncogene c-Myc through t(8;22) Translocation in Burkitt’s Lymphoma.” Oncogene 18 (9): 1745–53. https://doi.org/10.1038/sj.onc.1202468.

Hann, S. R., M. Dixit, R. C. Sears, and L. Sealy. 1994. “The Alternatively Initiated C-Myc Proteins Differentially Regulate Transcription through a Noncanonical DNA-Binding Site.” Genes & Development 8 (20): 2441–52. https://doi.org/10.1101/gad.8.20.2441.

Hann, Stephen R., Michael W. King, David L. Bentley, Carl W. Anderson, and Robert N. Eisenman. 1988. “A Non-AUG Translational Initiation in c-Myc Exon 1 Generates an N-Terminally Distinct Protein Whose Synthesis Is Disrupted in Burkitt’s Lymphomas.” Cell 52 (2): 185–95. https://doi.org/10.1016/0092-8674(88)90507-7.

Kovalchuk, Alexander L., Chen-Feng Qi, Ted A. Torrey, Lekidelu Taddesse-Heath, Lionel Feigenbaum, Sung Sup Park, Armin Gerbitz, et al. 2000. “Burkitt Lymphoma in the Mouse.” The Journal of Experimental Medicine 192 (8): 1183–90.

Laird-Offringa, I A, P Elfferich, H J Knaken, J de Ruiter, and A J van der Eb. 1989. “Analysis of Polyadenylation Site Usage of the C-Myc Oncogene.” Nucleic Acids Research 17 (16): 6499–6514.

Paronetto, Maria Paola, Ilaria Passacantilli, and Claudio Sette. 2016. “Alternative Splicing and Cell Survival: From Tissue Homeostasis to Disease.” Cell Death and Differentiation 23 (12): 1919–29. https://doi.org/10.1038/cdd.2016.91.

Spotts, G. D., S. V. Patel, Q. Xiao, and S. R. Hann. 1997. “Identification of Downstream-Initiated c-Myc Proteins Which Are Dominant-Negative Inhibitors of Transactivation by Full-Length c-Myc Proteins.” Molecular and Cellular Biology 17 (3): 1459–68. https://doi.org/10.1128/mcb.17.3.1459.

Thompson, E. B. 1998. “The Many Roles of C-Myc in Apoptosis.” Annual Review of Physiology 60: 575–600. https://doi.org/10.1146/annurev.physiol.60.1.575.

Wasylishen, Amanda R., Michelle Chan-Seng-Yue, Christina Bros, Dharmendra Dingar, William B. Tu, Manpreet Kalkat, Pak-Kei Chan, et al. 2013. “MYC Phosphorylation at Novel Regulatory Regions Suppresses Transforming Activity.” Cancer Research 73 (21): 6504–15. https://doi.org/10.1158/0008-5472.CAN-12-4063.

Watt, R., L. W. Stanton, K. B. Marcu, R. C. Gallo, C. M. Croce, and G. Rovera. 1983. “Nucleotide Sequence of Cloned CDNA of Human C-Myc Oncogene.” Nature 303 (5919): 725–28. https://doi.org/10.1038/303725a0.

Wierstra, Inken, and Jürgen Alves. 2008. “The C-Myc Promoter: Still MysterY and Challenge.” Advances in Cancer Research 99: 113–333. https://doi.org/10.1016/S0065-230X(07)99004-1.

Zaika, Alex I., Neda Slade, Susan H. Erster, Christine Sansome, Troy W. Joseph, Michael Pearl, Eva Chalas, and Ute M. Moll. 2002. “DeltaNp73, a Dominant-Negative Inhibitor of Wild-Type P53 and TAp73, Is up-Regulated in Human Tumors.” The Journal of Experimental Medicine 196 (6): 765–80. https://doi.org/10.1084/jem.20020179.

