## Supplemental data for "Alternative *c-MYC* mRNA transcripts as an additional tool for c-Myc2 and c-MycS production in BL60 tumors"

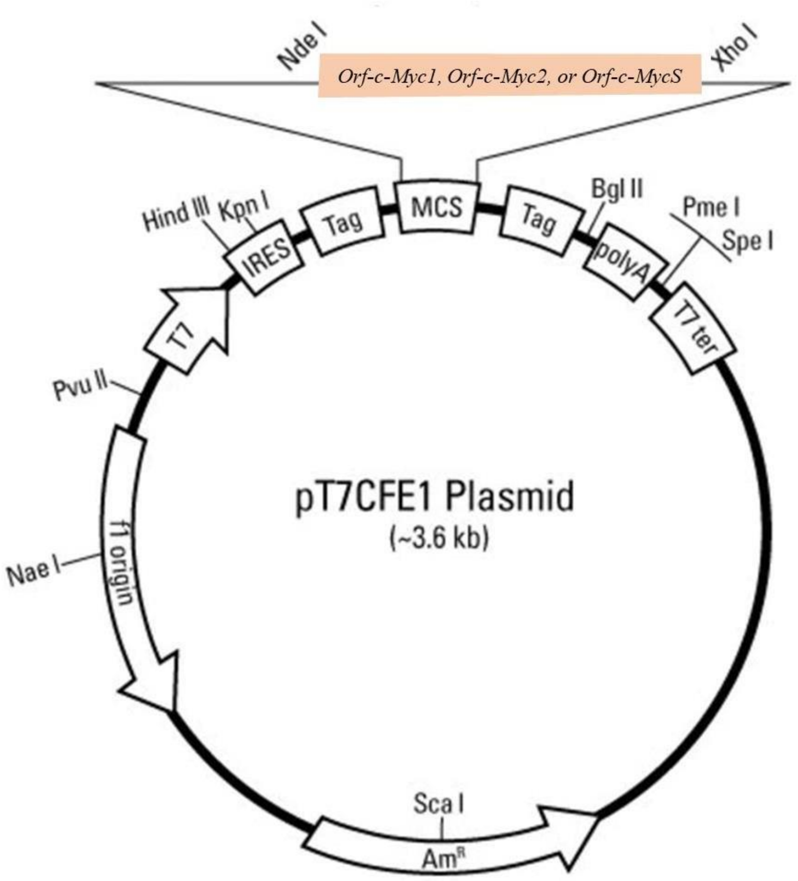


***<u>Figure S1:</u>*** ***Full-length open reading frames (Orf) encoding c-Myc1, c-Myc2 or c-MycS cloned into the pT7CFE1 vector subjected to transcription (T7 promoter) and translation using a mammalian in vitro translation system based on HeLa cell lysates.***

3’RACE

```
        P3’UTR1
        ---->
Genomic TAAGGAAAACGATTCCTTCTAACAGAAATGTCCTGAGCAATCACCTATGAACTTGTTTCAAATGCATGATCAAATGCAACCTCACAACCTTGGCTGAGTCTTGAGACTGAAAGATTTAGC 120
3’UTR1  TAAGGAAAACGATTCCTTCTAACAGAAATGTCCTGAGCAATCACCTATGAACTTGTTTCAAATGCATGATCAAATGCAACCTCACAACCTTGGCTGAGTCTTGAGACTGAAAGATTTAGC
3’UTR2  TAAGGAAAACGATTCCTTCTAACAGAAATGTCCTGAGCAATCACCTATGAACTTGTTTCAAATGCATGATCAAATGCAACCTCACAACCTTGGCTGAGTCTTGAGACTGAAAGATTTAGC

        -------------------------------------------------/continuation of 3’UTR/------------------------------------------------

Genomic TACACAATGTTTCTCTGTAAATATTGCCATTAAATGTAAATAACTTTAATAAAACGTTTATAGCAGTTACACAGAATTTCAATCCTAGTATATAGTACCTAGTATTATAGGTACTATAAA 360
3’UTR1  TACACAATGTTTCTCTGTAAATATTGCCATTAAATGTAAATAACTTTAATAAAACGTTTATAGCAGTTAAAAAAAAAAAAAAAAAAAAAAAAAAA
3’UTR2  TACACAATGTTTCTCTGTAAATATTGCCATTAAATGTAAATAACTTTAATAAAACGTTTATAGCAGTTACACAGAATTTCAATCCTAGTATATAGTACCTAGTATTATAGGTACTATAAA 360


Genomic CCCTAATTTTTTTTATTTAAGTACATTTTGCTTTTTAAAGTTGATTTTTTTCTATTGTTTTTAGAAAAAATAAAATAACTGGCAAATATATCATTGAGCCAAATCTTAAGTTGTGAATGT 480
3’UTR2  CCCTAATTTTTTTTATTTAAGTACATTTTGCTTTTTAAAGTTGATTTTTTTCTATTGTTTTTAGAAAAAATAAAATAACTGGCAAATATATCATTGAGCCAAATCTTAAAAAAAAAAAAA  ----> PolyA tail
```

**Figure S2: Nucleotide sequences of the identified 3'UTR types.** The translational TAA stop codon is indicated. Alternative AATAAA polyadenylation sites are underlined. P3’UTR1: Primer (Table1).

```
           P1: +1
            PFull1
Genomic     GACCCCCGAGCTGTGCTGCTCGCGGCCGCCACCGCCGGGCCCCGGCCGTCCCTGGCTCCCCTCCTGCCTCGAGAAGGGCAGGGCTTCTCAGAGGCTTGGCGGGAAAAAGAACGGAGGGAG
Trans1      GACCCCCGAGCTGTGCTGCTCGCGGCCGCCACCGCCGGGCCCCGGCCGTCCCTGGCTCCCCTCCTGCCTCGAGAAGGGCAGGGCTTCTCAGAGGCTTGGCGGGAAAAAGAACGGAGGGAG
Trans2      GACCCCCGAGCTGTGCTGCTCGCGGCCGCCACCGCCGGGCCCCGGCCGTCCCTGGCTCCCCTCCTGCCTCGAGAAGGGCAGGGCTTCTCAGAGGCTTGGCGGGAAAAAGAACGGAGGGAG

Genomic     GGATCGCGCTGAGTATAAAAGCCGGTTTTCGGGGCTTTATCTAACTCGCTGTAGTAATTCCAGCGAGAGGCAGAGGGAGCGAGCGGGCGGCCGGCTAGGGTGGAAGAGCCGGGCGAGCAG
Trans1      GGATCGCGCTGAGTATAAAAGCCGGTTTTCGGGGCTTTATCTAACTCGCTGTAGTAATTCCAGCGAGAGGCAGAGGGAGCGAGCGGGCGGCCGGCTAGGGTGGAAGAGCCGGGCGAGCAG
Trans2      GGATCGCGCTGAGTATAAAAGCCGGTTTTCGGGGCTTTATCTAACTCGCTGTAGTAATTCCAGCGAGAGGCAGAGGGAGCGAGCGGGCGGCCGGCTAGGGTGGAAGAGCCGGGCGAGCAG

Genomic     AGCTGCGCTGCGGGCGTCCTGGGAAGGGAGATCCGGAGCGAATAGGGGGGTTCGCCTCTGGCCCAGACCTCCCGCTGATGCCCCAGCCAGCGGTCCGCAACCCTTGCCGCATCCACGAAA
Trans1      AGCTGCGCTGCGGGCGTCCTGGGAAGGGAGATCCGGAGCGAATAGGGGGGTTCGCCTCTGGCCCAGACCTCCCGCTGATGCCCCAGCCAGCGGTCCGCAACCCTTGCCGCATCCACGAAA
Trans2      AGCTGCGCTGCGGGCGTCCTGGGAAGGGAGATCCGGAGCGAATAGGGGGGTTCGCCTCTGGCCCAGACCTCCCGCTGATGCCCCAGCCAGCGGTCCGCAACCCTTGCCGCATCCACGAAA

Genomic     CTTTGCCCATAGCAGCGGGCGGGCTCTTTGCACTGGAACTTACAACACCCGAGTAAGGACGCGACTCTCCCGACGCGGGGAGGCTATTCTGCCCATTTGGGGACACTTCCCCGCCGCTGC
Trans1      CTTTGCCCATAGCAGCGGGCGGGCTCTTTGCACTGGAACTTACAACACCCGAGTAAGGACGCGACTCTCCCGACGCGGGGAGGCTATTCTGCCCATTTGGGGACACTTCCCCGCCGCTGC
Trans2      CTTTGCCCATAGCAGCGGGCGGGCTCTTTGCACTGGAACTTACAACACCCGAGTAAGGACGCGACTCTCCCGACGCGGGGAGGCTATTCTGCCCATTTGGGGACACTTCCCCGCCGCTGC
                                           PvuII           (c-Myc1)
Genomic     CAGGACCCGCTTCTCTGAAAGGCTCTCCTTGCAGCTGCTTAGACGCTGGATTTTTTTCGGGTAGTGGAAAACCAGGTAAGCACCGAAGTCCACTTGCCTTTTAATTTATTTTTTTATCAC
Trans1      CAGGACCCGCTTCTCTGAAAGGCTCTCCTTGCACCTGCTTAGACGCTGGATTTTTTTCGGGTAGTGGAAAACCAG------------intron1-----------------------

Trans2      CAGGACCCGCTTCTCTGAAAGGCTCTCCTTGCAGCTGCTTAGACGCTGGATTTTTTTCGGGTAGTGGAAAACCAG------------intron1-----------------------
            ----------------------------//continuation of intron1//--exon2------------------------------------------------------
                                                                                  (c-Myc2)
Genomic     CC----AAGACCCCTTTAACTCAAGACTGCCTCCCGCTTTGTGTGCCCCGCTCCAGCAGCCTCCCGCGACGATGCCCCTCAACGTTAGCTTCACCAACAGGAACTATGACCTCGACTACG
Trans1                                                              CAGCCTCCCGCGACGATGCCCCTCAACGTTAGCTTCACCAACAGGAACTATGACCTCGACTACG
                                                                                   M  P  L  N  V  S  F  T  N  R  N  Y  D  L  D  Y
Trans2                                                                                                             GAACTATGACCTCGACTACG

            ------------------------------//Continuation of exon2//-------------------------------------------------------------
                                                                                   (c-MycS)
Genomic     TCACACCCTTCTCCCTTCGGGGAGACAACGACGGCGGTGGCGGGAGCTTCTCCACGGCCGACCAGCTGGAGATGGTGACCGAGCTGCTGGGAGGAGACATGGTGAACCAGAGTTTCATCT
Trans1      TCACACCCTTCTCCCTTCGGGGAGACAACGACGGCGGTGGCGGGAGCTTCTCCACGGCCGACCAGCTGGAGATGGTGACCGAGCTGCTGGGAGGAGACATGGTGAACCAGAGTTTCATCT
              T  P  F  S  L  R  G  D  N  D  G  G  G  G  S  F  S  T  A  D  Q  L  E  M  V  T  E  L  L  G  G  D  M  V  N  Q  S  F  I   ----> c-Myc2
Trans2      TCACACCCTTCTCCCTTCGGGGAGACAACGACGGCGGTGGCGGGAGCTTCTCCACGGCCGACCAGCTGGAGATGGTGACCGAGCTGCTGGGAGGAGACATGGTGAACCAGAGTTTCATCT
                                                                                   M  V  T  E  L  L  G  G  D  M  V  N  Q  S  F  I   ----> c-MycS
```

***Figure S3:*** ***Nucleotide sequences of Trans1 and Trans2 transcripts compared to the corresponding genomic structure.*** *Underlined splice donor (GT), acceptor (AG) sites, and initiation codons (CTG, ATG) are indicated. The first amino acids of c-Myc2 and c-MycS are shown. Translation from CTG of Trans1 is prevented by the PvuII mutation. +1: Transcription initiation site from the P1 promoter. PFull1: Primer (Table1); M: The first amino acid, methionine. The rest of c-MYC sequence until the translation stop codon (TAA) is not shown.*

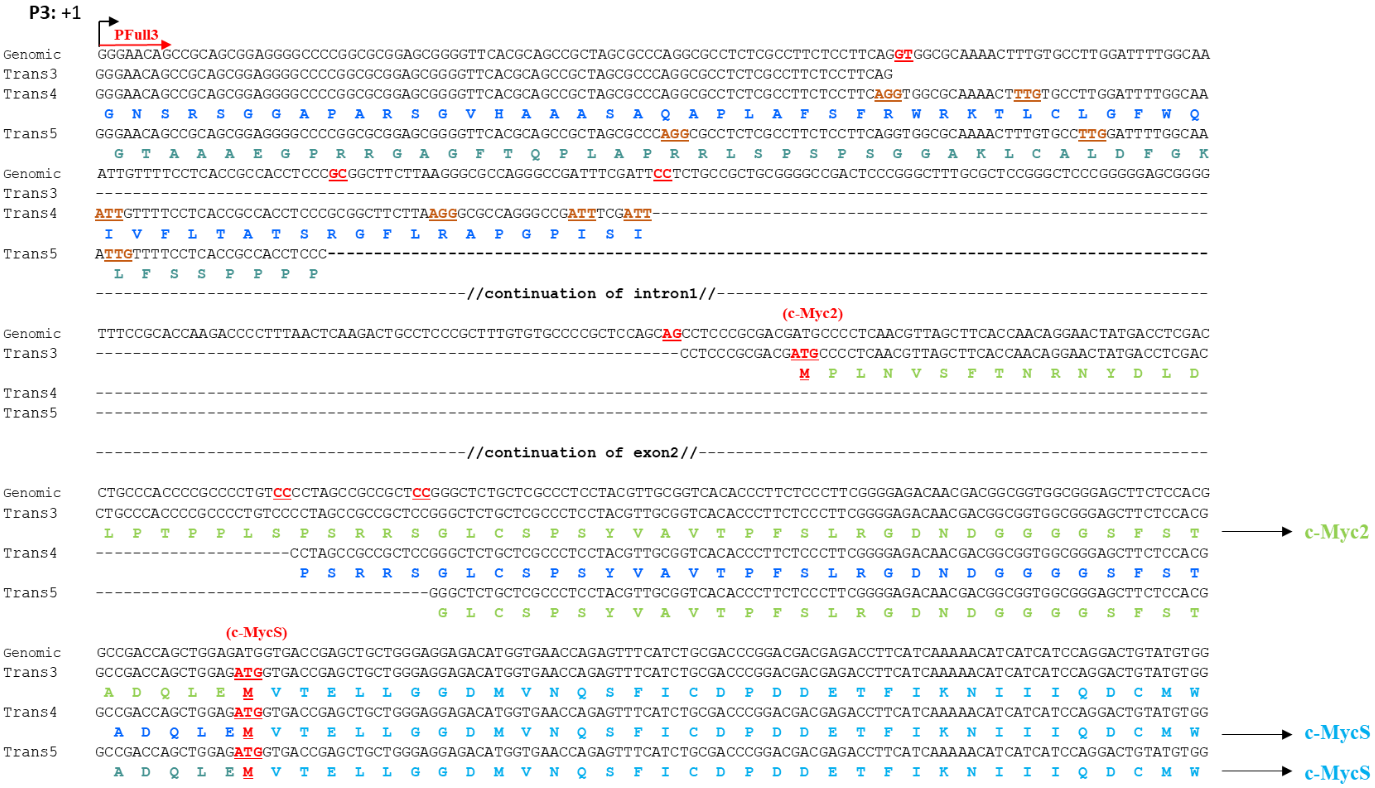


***<u>Figure S4:</u>* Nucleotide sequences of Trans3, Trans4, and Trans5 transcripts compared to the corresponding genomic structure.** *Underlined dinucleotides represent the splice sites found. The initiation codons for c-Myc2 and c-MycS are indicated. Trans3 translation predicts c-Myc2 protein. Curiously and despite different splicings, Trans4 and Trans5 transcripts do not have a stop codon in their 5'UTR region. Potential non-ATG initiation sites are highlighted in brown. +1: transcription initiation site from the P3 promoter. PFull3: primer (Table1); M: The first amino acid, methionine. The rest of c-MYC sequence until the translation stop codon (TAA) is not shown.*

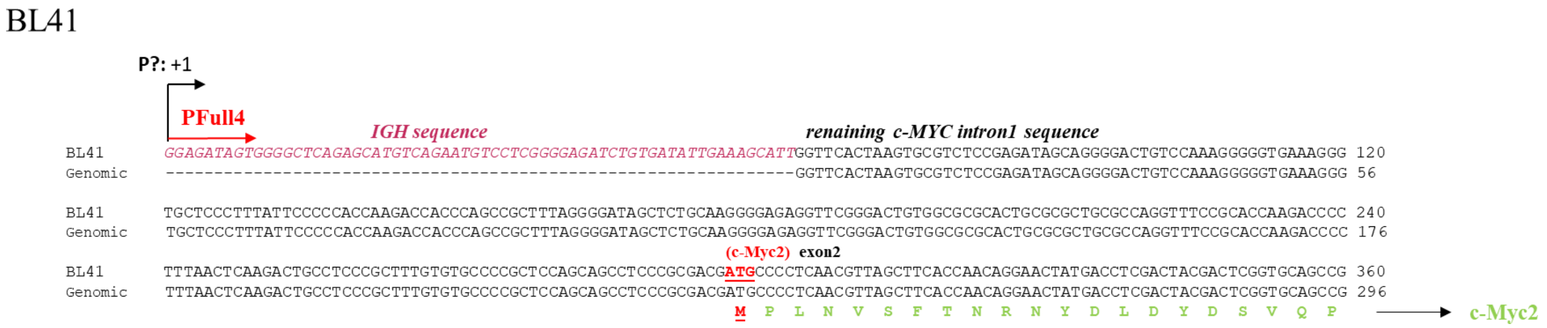


***<u>Figure S5:</u>* Nucleotide sequences of the only transcript expressed from a yet to be determined promoter (P?).**

*The transcript exhibits an IGH sequence. This transcript encodes c-Myc2. PFull4: primer (Table1); M: methionine. The rest of c-MYC sequence until the translation stop codon (TAA) is not shown.*
